# Social fluidity mobilizes contagion in human and animal populations

**DOI:** 10.1101/170266

**Authors:** Ewan Colman, Vittoria Colizza, Ephraim M. Hanks, David P. Hughes, Shweta Bansal

## Abstract

Humans and other group-living animals tend to distribute their social effort disproportionately. Individuals predominantly interact with a small number of close companions while maintaining weaker social bonds with less familiar group members. By incorporating this behaviour into a mathematical model we find that a single parameter, which we refer to as social fluidity, controls the rate of social mixing within the group. We compare the social fluidity of 13 species by applying the model to empirical human and animal social interaction data. To investigate how social behavior influences the likelihood of an epidemic outbreak we derive an analytical expression of the relationship between social fluidity and the basic reproductive number of an infectious disease. For highly fluid social behaviour disease transmission is revealed to be density-dependent. For species that form more stable social bonds, the model describes frequency-dependent transmission that is sensitive to changes in social fluidity.

---

**S**ocial behavior is fundamental to the survival of many species. It allows the formation of social groups providing fitness advantages from greater access to resources and better protection from predators [1]. Structure within these groups can be found in the way individuals communicate across space, cooperate in sexual or parental behavior, or clash in territorial or mating conflicts [2]. While animal societies are usually studied independently of each other, studying their differences has potential to reveal new insights into the nature of social living [3, 4].

When social interaction requires shared physical space it can also be a conduit for the transmission of infectious disease [5]. In a typical infectious disease model, if the disease spreads through the environment then the transmission rate is assumed to scale proportionally to the local population density [6, 7]. Alternatively, if transmission requires close proximity encounters that only occur between bonded individuals, then we expect social connectivity to determine the outcome. These two paradigms are known in the literature as density-dependence and frequency-dependence [8].

The problem, however, is that real diseases are not so easy to categorize [9]. For example, as social groups grow in size, new bonds must be created to maintain cohesiveness [10]. To manage the time and cognitive effort required to create these bonds, individuals tend to interact mostly with a small number of close companions while maintaining cohesion with the wider group through less frequent contact [11–13]. For an infectious disease, this creates fewer transmission opportunities than we would expect to see in a group with highly fluid social dynamics. The extent to which group size amplifies the transmission rate therefore depends on how individuals choose to distribute their social effort between strong and weak ties [14].

The purpose of this study is to address two questions. Firstly, can we quantify the variability in how individuals choose to distribute their social effort within a group, and secondly, what will this tell us about the effect that population density has on disease transmission?

There is growing evidence for the disproportionate distribution of social effort in human telecommunication [15–18]. Quantifying this aspect of sociality in animal systems, however, has been held back by the limitations of the data. One challenge, which we address here, is the bias introduced by variation in activity levels across the social group [19]. Additionally, while heterogeneous interaction frequencies and temporal dynamics have become common in epidemiological models [20, 21], little has been done to incorporate the variability in how the individual chooses to expend their social effort.

In the first part of this paper we introduce a mathematical model founded on the concept of *social fluidity* which we define as variability in the amount of social effort the individual invests in each member of their social group. Using openly available data, we estimate the social fluidity of 57 human and animal social systems. In the second part we derive an expression for the basic reproductive number of an infectious disease in the social fluidity model and demonstrate its accurately in predicting simulated outcomes. Furthermore, social fluidity emerges as a coherent mathematical framework providing the smooth connection between density-dependent and frequency-dependent disease systems.

## Characterizing social behaviour

Our first objective is to measure social behaviour in a range of human and animal populations. We start by introducing a model that captures a hidden element of social dynamics: how individual group members distribute their social effort. We mathematically describe the relationships between social variables that are routinely found in studies of animal behavior, the number of social ties and the number of interactions observed, and apply the model to empirical data to reveal behavioural differences between several species.

### Social behavior model

Consider a closed system of *N* individuals and a set of interactions between pairs of individuals that were recorded during some observation period. These observations can be represented as a network: each individual, *i*, is a *node*; an *edge* exists between two nodes *i* and *j* if at least one interaction was observed between them; the *edge weight, w*_*i,j*_, denotes the number of times this interaction was observed. The total number of interactions of *i* is denoted *strength, s*_*i*_ = Σ_*j*_ *w*_*i,j*_, and the number of nodes with whom *i* is observed interacting is its *degree, k*_*i*_ [22].

We define *x*_*j*|*i*_ to be the probability that an interaction involving *i* will also involve node *j*. Therefore the probability that at least one of these interactions is with *j* is 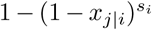. The main assumption of the model is that the values of *x*_*j*|*i*_ over all *i, j* pairs are distributed according to a probability distribution, *ρ*(*x*).^1^ Thus, if a node interacts *s* times, the marginal probability that an edge exists between that node and any other given node in the network is

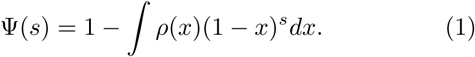

Our goal is to find a form of *ρ* that accurately reproduces network structure observed in real social systems. Motivated by our exploration of empirical interaction patterns from a variety of species (Fig. S1), we propose that *ρ* has a power-law form:

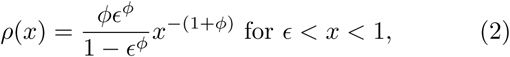

where *ϕ* (> 0) controls the variability in the values of *x*, and *ϵ* simply truncates the distribution to avoid divergence. Combining (1) and (2) we find

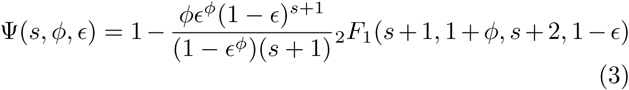

where the notation _2_*F*_1_ refers to the Gauss hypergeometric function [23]. It follows from Σ_*j*_ *x*_*j*|*i*_ = 1 that

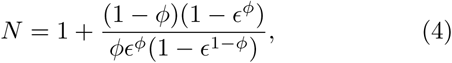

which can be solved numerically to find ϵ for given values of *N* and *ϕ*. The expectation of the degree is *κ*(*s, ϕ, N*) = (*N* 1)Ψ(*s, ϕ, ϵ*).

Fig. 1 illustrates how the value of *ϕ* can produce different types of social behavior. As *ϕ* is the main determinant of social behaviour in our model, we use the term *social fluidity* to refer to this quantity. Low social fluidity (*ϕ* ≪ 1) produces what we might describe as “allegiant” behavior: interactions with the same partner are frequently repeated at the expense of interactions with unfamiliar individuals. As *ϕ* increases, the model produces more “gregarious” behavior: interactions are repeated less frequently and the number of partners grows faster. While names like “social strategy” and “loyalty” have been applied to similar concepts [24, 25], fluidity, as a property of matter, is a useful metaphor for communicating the main idea behind this model.

**Figure 1:**
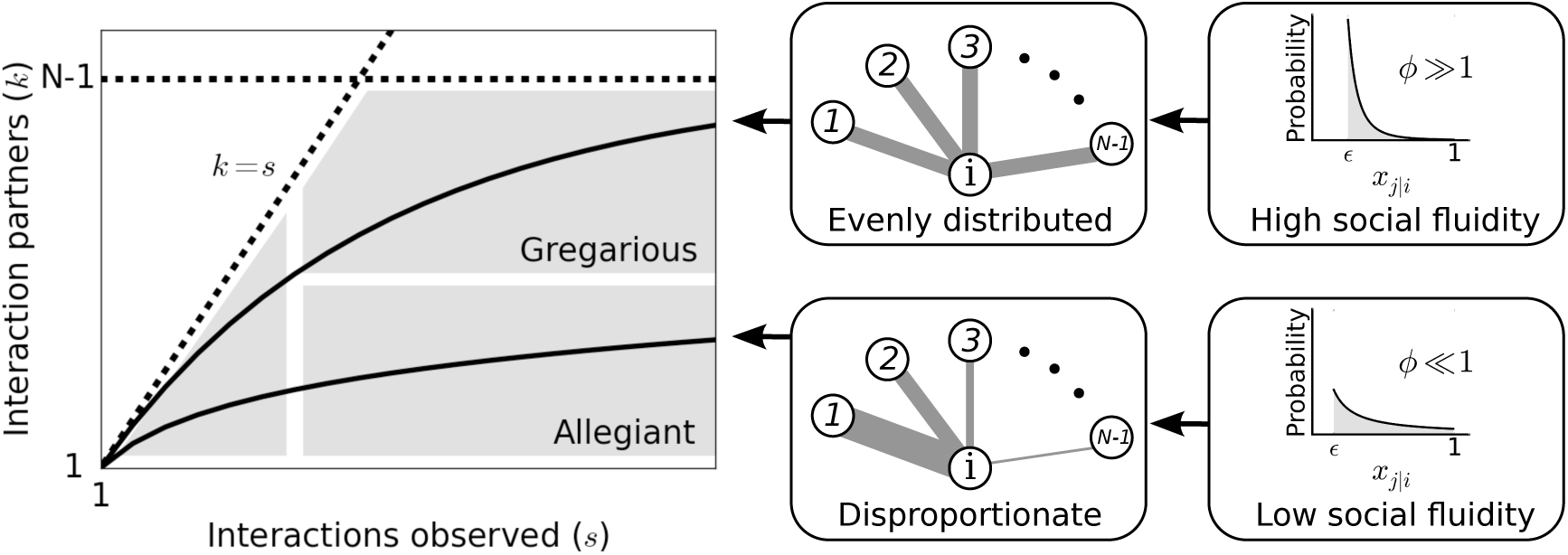
Left: Each individual can be represented as a single point on this plot. Dashed lines mark the boundary of the region where data points can feasibly be found. The mean degree is plotted for two values of *ϕ* representing two possible types of social behavior; as the number of observed interactions grows, the set of social contacts increases; the rate at which it increases influences how we categorize their social behavior. Middle: The weight of the edges between *i* and the other nodes represents the propensity of *i* to interact with each of the other individuals in the group. Right: Probability distributions that correspond to the different levels of evenness in the contact propensities, both distributions are expressed by Eq.(2).

#### Estimating social fluidity in empirical networks

To understand the results of the model in the context of real systems we estimate *ϕ* in 57 networks from 20 studies of human and animal social behavior (further details in the supplement) [26–46], focusing our attention to those interactions which are capable of disease transmission (i.e. those that, at the least, require close spatial proximity).

Each dataset provides the number of interactions that were observed between pairs of individuals. We assume that the system is closed, and that the total network size (*N*) is equal to the number of individuals observed in at least one interaction. To estimate social fluidity we find the value of *ϕ* that minimizes Σ_*i*_[*k*_*i*_−*κ*(*s*_*i*_, *ϕ, N*)]^2^ (the total squared squared error between the observed degrees and their expectation given by the model). Being estimated from the relationship between strength and degree, and not their absolute values, social fluidity is a good candidate for comparing social behavior across different systems as it is independent of the distributions of *s*_*i*_ or *k*_*i*_, and of the timescale of interactions.

Fig. 2 shows the estimated values of *ϕ* for all networks in our study. We organize the measurements of social fluidity by interaction type. Aggressive interactions have the highest fluidity (which implies that most interactions are rarely repeated between the same individuals), while grooming and other forms of social bonding have the lowest (which implies frequent repeated interactions between the same individuals). Social fluidity also appears to be related to species: ant systems cluster around *ϕ* = 1, monkeys around *ϕ* = 0.5, humans take a range of values that depend on the social environment. Sociality type does not appear to affect *ϕ*; sheep, bison, and cattle have different social fluidity compared to kangaroos and bats, though they are all cat-egorized as fissionfusion species [3].

**Figure 2:**
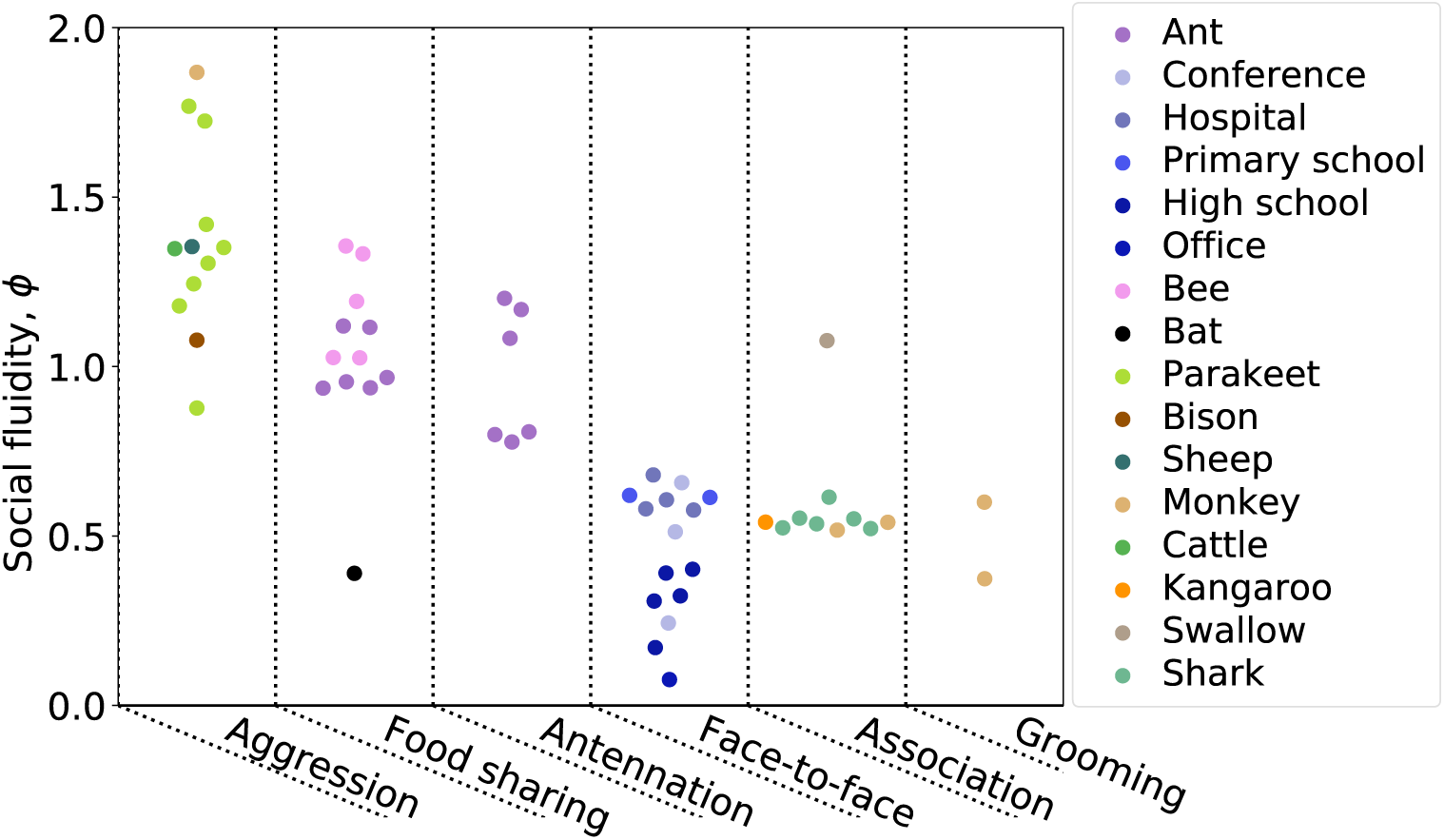
Each point represents a human or animal system for which social fluidity was estimated. Results are organized by interaction type: aggression includes fighting and displays of dominance, food sharing refers to mouth-to-mouth passing of food, antennation is when the antenna of one insect touches any part of another, space sharing interactions occur with spatial proximity during foraging, face-to-face refers to close proximity interactions that require individuals to be facing each other, association is defined as co-membership of the same social group.

There is no significant correlation between the mean number of interactions per individual 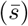 and social flu-idity (Pearson *r*^2^ = 0.02, *p* = 0.26), which implies that sampling bias does not affect the estimation of social fluidity. Similarly, network size does not correlate with *ϕ* (Pearson *r*^2^ = 0.02, *p* = 0.33). Larger values of *ϕ* correspond to higher mean degrees (Pearson *r*^2^ = 0.27, *p* < 0.001) and lower variability in the distribution of edge weights (measured as the index of dispersion of *w*_*i,j*_; Pearson *r*^2^ = 0.26, *p* < 0.001). Weight variability and mean degree are uncorrelated in these data (Pearson *r*^2^ = 0.01, *p* = 0.59) implying that *ϕ* combines these two entirely distinct features of social behavior.

Finally, the modularity of the network (computed by the Louvain method on the unweighted network [47]) is negatively correlated with *ϕ* (*r*^2^ = 0.57, *p* < 0.001). This is expected as individuals tend to be loyal to those within the same module while maintaining weaker connections with the remaining network in all but one network the mean weight of edges within modules is higher than the mean weight of edges between modules (supplementary document).

### Characterizing disease spread with social fluidity

Our objective is to characterize how social behavior influences the susceptibility of the group to infectious disease in a range of human and animal social systems. Intuitively, we expect an infected individual in a group with low social fluidity to expose fewer susceptible group members to the pathogen than they would in a group with highly fluid social dynamics. We explore this idea by introducing a analytical transmission model that incorporates social fluidity. Using this model, we mathematically characterize the impact of social fluidity on density dependence, and apply the model to empirical networks to predict disease spread.

#### Disease transmission model

We consider the transmission of an infectious disease on the social behavior model introduced in the previous section. An infectious node *i* interacting with a susceptible node *j* will transmit the infection with probability *β*. The node will recover from infection with rate *γ*, assuming an exponential distribution of the length of the infectious period. The probability that the infection is transmitted from *i* to any given *j* is

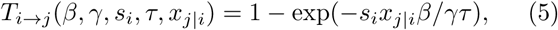

assuming that the interactions *s*_*i*_ of *i* are distributed randomly across an observation period of duration *τ*.

By integrating Eq. (5) over all possible values *x*_*j*|*i*_ and and infectious period durations and multiplying by the number of susceptible individuals (*N –* 1) we obtain the expected number of infections caused by individual *i*,

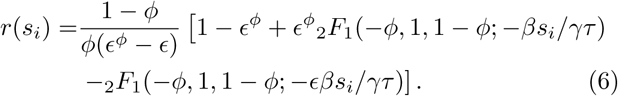

The basic reproductive number (usually denoted *R*_0_) is defined as the mean number of secondary infections caused by a typical infectious individual in an otherwise susceptible population [48]. We will use the notation 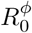 to signify the *social fluidity reproductive number*, that is the analogue of *R*_0_ derived from our social behaviour model.

We assess the relation of the reproductive number with the population density by focusing on a special case where every node has the same strength, i.e *s*_*i*_ = *s* for all *i*, so that 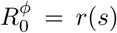. Furthermore, we choose 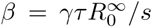 where 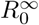 is 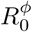 as *ϕ* → *∞*, i.e, a constant that represents what the basic reproductive number would be if every new interaction occurred between a pair of individuals who have not previously interacted with each other.

Fig. 3 shows the effect of social fluidity on the density dependence of the disease. At small population sizes, 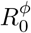 increases with *N* and converges as *N* goes to ∞ (Fig. 3**A**). The rate of this convergence increases with *ϕ*, and the limit it converges to is higher, meaning that *ϕ* determines the extent to which density affects the spread of disease. As *N* → *∞*, we find that 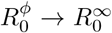 for *ϕ* > 1. When *ϕ* < 1, 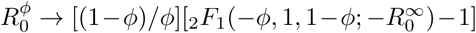. At these values of *ϕ* the disease is constrained by individuals choosing to repeat interactions despite having the choice of infinitely many potential interaction partners (Fig 3**B**).

**Figure 3:**
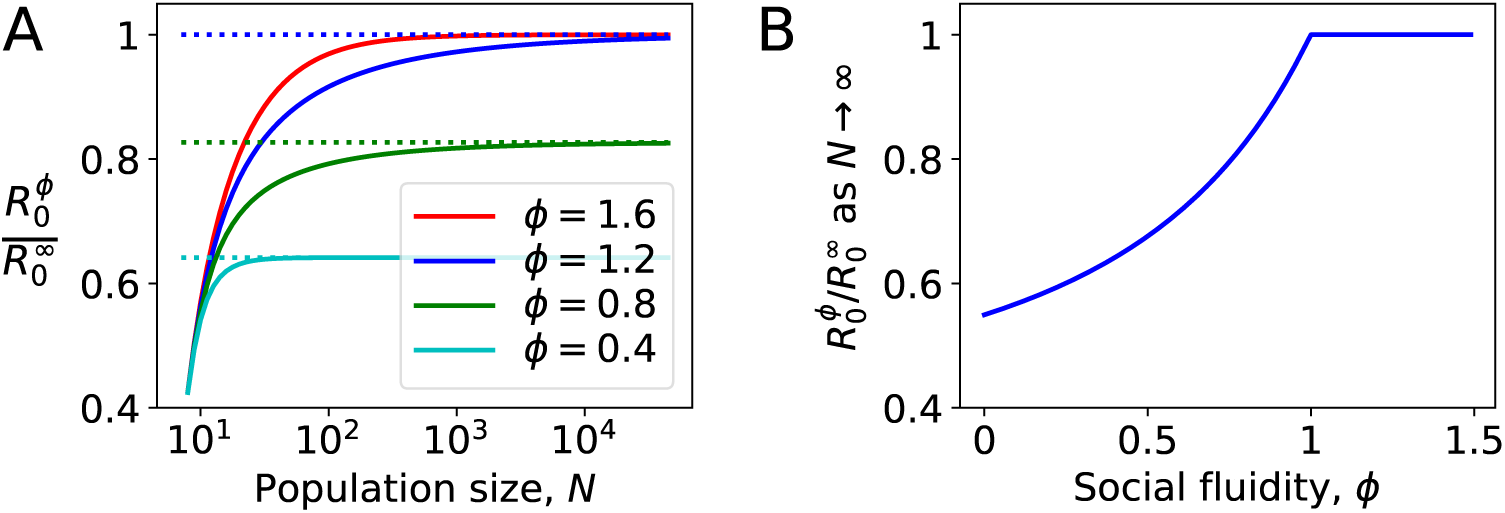
Density dependence in populations where every node has the same strength. **A:** For different values of social fluidity, *ϕ*, we show 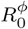 (from Eq.(6)) as a function of *N* (from Eq.(4)) through their parametric relation with ϵ. Dashed lines show the limit for large *N*. B: In large populations 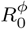 increases with *ϕ* up to *ϕ* = 1. Beyond this value, infections occur as frequently as they would if every new interaction occurs between a pair of individuals who have not previously interacted with each other.

#### Estimating infection spread in empirical networks with heterogeneous connectivity

To apply this analogue of a reproductive number to an animal-disease system, we need to account for heterogeneous levels of social connectivity in the given population and thus the tendency for infected individuals to be those with a greater number of social partners [49]. For the basic reproductive number, this is often done using the mean *excess degree*, i.e. the degree of an individual selected with probability proportional to their degree [50]. Following a similar reasoning, we define 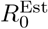, which incorporates the effect of social fluidity, as the expected number of infections (*r*(*s*_*i*_)) caused by an individual that has been selected with probability proportional to their degree (*k*_*i*_):

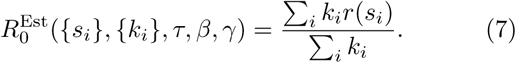

Given the degree and strength of each individual in a network, the duration over which those interactions occurrred, and the transmission and recovery rates of the disease, we are able to estimate *ϕ*, compute Eq.(6) for each individual, and finally use Eq.(7) to derive a statistic that provides a measure of the risk of the host population to disease outbreak.

#### Numerical validation using empirical networks

We simulated the spread of disease through the interactions that occurred in the empirical data (materials and methods). We compute 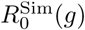, defined as the ratio of the number of individuals infected at the (*g* + 1)-th generation to the number infected at the *g*-th generation over 10^3^ simulated outbreaks, for *g* = 0, 1, 2 (*g* = 0 refers to the initial seed of the outbreak).

Table 1 shows the Pearson correlation coefficient between 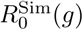 and its corresponding value 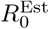 obtained Eq.(7). For comparison, the correlation is shown for other indicators and network statistics. The results correspond to one set of simulation conditions, and are robust across a wide range of parameter combinations (see supplementary tables). Note that a different value of *β* was chosen for each network to control for the varying interaction rates between networks while keeping the upper bound 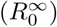 constant (materials and methods). Thus, the mean strength does not have a significant effect on 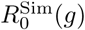.

**Table 1:** The Pearson correlation coefficient between quantities calculated on the network and the simulated disease outcomes (with 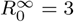). Results that are significant with *p* < 0.01 are labelled with *.

| | Corr. with $R_0^{\text{Sim}}(g = 1)$ |
| --- | --- |
| $R_0^{\text{Est}}$ | 0.91* |
| Social fluidity | 0.73* |
| Excess degree | 0.64* |
| Mean degree | 0.53* |
| Network size | 0.47* |
| Mean strength | -0.07 |
| Mean clustering | -0.15 |
| Mean edge weight | -0.45* |
| Edge weight variability | -0.48* |
| Modularity | -0.60* |

These correlations support a known result regarding repeat contacts in network models of disease spread: that indicators of disease risk that are derived solely from the degree distribution are unreliable and the role of edge weights should not be neglected [51,52]. After transmission has occurred from one individual to another, repeating the same interaction serves no advantage for disease (most directly-transmitted microparasites are not dose-dependent). Since a large edge weight implies a high frequency of repeated interactions, networks with a higher mean weight tend to have lower basic reproductive numbers. Furthermore, variability in the distribution of weights concentrates a yet larger proportion of interactions onto a small number of edges, further increasing the number of repeat interactions and reducing the reproductive number.

Correlation between modularity and 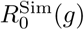 is partly due to the strong correlation between modular networks and those with high social fluidity. Consistent with other evidence [53], this suggests that transmission events occur mostly within the module of the seed node, with weaker social ties facilitating transmission to other modules. The effect of clustering (a measure of the number of connected triples in network [54]) correlates with smaller 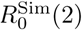, consistent with other theoretical work [51, 55].

Finally, we find the model estimate of the social fluidity reproductive number 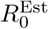 to be, on average, within 10% of the simulated value, 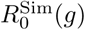. At *g* = 2 the amount of error is larger (to up to 29% for some parameter choices). Prediction accuracy at this generation is negatively correlated with the mean clustering coefficient. This is not surprising as 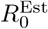 does not account for the accelerated depletion of susceptible neighbours that is known to occur in clustered networks [51, 55]. No other properties of the network affect the accuracy of 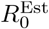 consistently across all parameter combinations (see supplementary tables).

## Discussion

We proposed a measure of fluidity in social behavior which quantifies how much mixing exists within the social relationships of a population. While social networks can be measured with a variety of metrics including size, connectivity, contact heterogeneity and frequency, our methodology reduces all such factors to a single quantity allowing comparisons across a range of human and animal social systems. Social fluidity correlates with both the density of social ties (mean degree) and the variability in the weight of those ties, though these quantities do not correlate with each other. Social fluidity is thus able to combine these two aspects seamlessly in one quantity.

By measuring social fluidity across a range of human and animal systems we are able to rank social behaviors. We identify aggressive interactions as the most socially fluid; this indicates a possible learning effect whereby each aggressive encounter is followed by a period during which individuals avoid further aggression with each other [56]. At the opposite end of the scale, we find interactions that strengthen bonds (and thus require repeated interactions) such as grooming in monkeys [57] and food-sharing in bats [33]. The fact that food-sharing ants are far more fluid than bats, despite performing the same kind of interaction, reflects their eusocial nature and the absence of any need to consistently reinforce bonds with their kin [58].

Most studies that aim to describe and quantify social structure are met with a number of challenges, including ours. First, the degree of an individual, for example, is known to scale with the length of the observation period [59]. By focusing not on the absolute value of degree, but instead on how degree scales with the number of observations, our analysis controls for this bias. Second, observed interactions have been assumed to persist over time [60]. In our model, only the distribution of edge weights remains constant through time, an assumption consistent with growing evidence [25, 61]. Third, duration of contacts is known to be important for disease spread [52]. We did not include explicitly the duration of each contact in our model, since this information was only available in a fraction of the datasets [62]. There is therefore potential to improve the applicability of this model as more high resolution data becomes openly available.

Our estimate of reproductive number derived from social fluidity provides a better predictor for the epidemic risk of a host population, going beyond predictors based on density or degree only. To illustrate this point, the social network of individuals at a conference (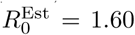; `conference_0`, supplementary document) is predicted to be at higher risk compared to the social network at a school (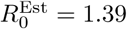; `highschool_0`), despite having a smaller size and lower connectivity (*N* = 93 vs. *N* = 312, and 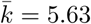 vs. 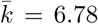, respectively). The discrepancy in the risk prediction comes from the lower frequency of repeated contacts between individuals in the conference, compared to the school. Interactions between infectious individuals and those they have previously infected are redundant in terms of transmission. This dynamic is nicely captured by the social fluidity, with *ϕ* = 0.66 for the conference and *ϕ* = 0.40 for the high school.

Unlike previous work that explores the disease consequences of population mixing [63, 64], our analysis allows us to investigate this relation across a range of social systems. We see, for example, how the relationship between mixing and disease risk scales with population density. For social systems that have high values of social fluidity, 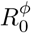 is highly sensitive to changes in *N*, whereas this sensitivity is not present at low values of *ϕ*. This corroborates past work on the scaling of transmission being associated to heterogeneity in contact [65,66]. Going beyond previous work, our model captures in a coherent theoretical framework both density-dependence and frequency-dependence, and social fluidity is the measure to tune from one to the other in a continuous way. Since many empirical studies support a transmission function that is somewhere between these two modeling paradigms [7, 67–69], the modeling approaches applied in this paper can be carried forward to inform transmission relationships in future disease studies.

## Materials & Methods

### A. Python libraries

Mean clustering coefficients were computed using the *networkx* Python library. To evaluate the hypergeometric function in (3) we used the *hyp2f1* function from the *scipy*.*special* Python library. Numerical solutions to Eq.(4) using the *fsolve* function from the *scipy*.*optimize* Python library. All scripts, data, and documentation used in this study are available through https://github.com/EwanColman/Social-Fluidity.

### B. Data handling

Only freely available downloadable sources of data have been used for this study. Details of the experimentation and data collection can be found through their respective publications. Here we note some additional processes we have applied for our study.

Each human contact dataset lists the identities of the people in contact, as well as the 20-second interval of detection [26–29, 32]. Any sequence of consecutive time intervals for which contact is detected between two individuals is considered to be one interaction. To exclude contacts detected while participants momentarily walked past one another, only contacts detected in at least two consecutive intervals are considered interactions. Data were then separated into 24 hour subsets.

Bee trophallaxis provided experimental data for 5 unrelated colonies under continuous observation. We use the first hour of recorded data for each colony [46]. The ant trophallaxis study provided 6 networks: 3 unrelated colonies continuously observed under 2 different experimental conditions [30]. Ant antennation study provided 6 networks: 3 colonies, each observed in 2 sessions separated by a two week period. The bat study collected individual data at different times and under different experimental conditions [33]. For bats that were studied on more than one occasion we use only the first day they were observed.

Some data sets provided data for group membership collected through intermittent, rather than continuous, observation [34–38]. We construct networks from these data by recording an interaction when two individuals were seen to be in the same group during one round of observation. The shark data was divided into 6 datasets, each one constructed from 10 consecutive observations, and spread out through the full time period over which the data was collected.

For the grooming data [39, 40], if one animal was grooming another during one round of observations then this would be recorded as a directed interaction. Similarly for aggressive interactions [41–45, 56]. When an animal was determined to be the winner of a dominance encounter then this would be recorded as a directed interaction between the winner and the loser. We consider interaction in either direction to be a contact in the network.

We considered including two rodent datasets in which interaction is defined as being observed within the same territorial space [67,69]. We did not find this suitable for our analysis since the network we obtain, and the consequent results are sensitive to setting of arbitrary threshold values regarding what should, or should not, be considered sufficient contact for an interaction.

For data that did not contain the time of each interaction, contact time series were generated synthetically. For those datasets, the interactions between each pair were given synthetic timestamps in three different ways, Poisson: the time of each interaction is chosen uniformly at random from {0, 1, …, 10^4^} seconds, Circadian: chosen uniformly at random from {0, 1, …, 3333, 6666, …, 10^4^}, and Bursty: interaction times occur with power-law distributed inter-event times adjusted to give an expected total duration of 10^4^ seconds.

### C. Disease simulation

Simulations of disease spread were executed using the contacts provided by the datasets. The the bat network was omitted from this part since these data were collected over a series of independent experiments carried out at different times and under different experimental treatments.

In one run of the simulation, one seed node is randomly chosen from the network and, at a randomly selected point in time during the duration of the data, transitions to the infectious state. The duration for which they remain infectious is a random variable drawn from an exponential distribution with mean 1*/γ*. During this time any contact they have with other individuals who have not previously been infected will cause an infection with probability *β*.

The simulation runs until all individuals who were infected at the second generation of the disease, i.e. those infected by those infected by the seed, have recovered. The datasets are ‘looped’ to ensure that the timeframe of the data collection does not influence the outcome. In other words, immediately after the latest interaction, the interactions are repeated exactly as they were originally. This continues to happen until the termination criteria is met.

We set the parameters to normalise for the variation in contacts rates between networks. To achieve this we consider a hypothetical counterpart to each network in which the strength of every node is the same, but each interaction occurs between a pair of individuals who have not previously interacted. This is equivalent to *ϕ* → *∞*. Under these conditions *x*_*j*|*i*_ = 1*/*(*N* − 1) for all pairs *i, j*. It follows that Eq. (5) becomes *T_i_*→*_j_ ≈ siβ/γτ* (*N* − 1), then *r*(*s_i_*) *≈ siβ/γτ*, and, since *k_i_* = *s_i_* for all nodes *i*, Eq. (7) gives

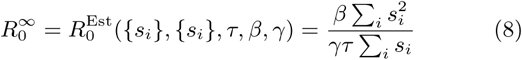

The value of 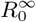 can be chosen arbitrarily. Then, by setting *γ* = 1*/τ* and 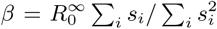 we guarantee that Eq. (8) holds for every network. To test that our results hold over a range of disease scenarios we repeat our analysis with 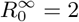, 3, and 4.

## Supporting information

Supplementary results

## Acknowledgments

This work was supported by NSF grant number 1414296. We are grateful for insightful feedback from Pratha Sah. We also thank all the researchers who have made their behavioral data openly accessible, making this study possible.

## Footnotes

1 *x*_*j*|*i*_ are subject to network interdependencies. Specifically, *AX* = *X*^*T*^ *A* and *X***1** = **0**, where *X* is a matrix whose *i, j* entry is –1 if *i* = *j* and *x*_*j*|*i*_ otherwise, *A* is any diagonal matrix with positive entries, and **0** and **1** are column vectors of length *N* containing only 0 and 1, respectively. Thus, *ρ*(*x*) is the distribution of marginal *x*_*j*|*i*_ values of the joint distribution *P*(*X*).

